# SARS-CoV-2 Omicron Variant: ACE2 Binding, Cryo-EM Structure of Spike Protein-ACE2 Complex and Antibody Evasion

**DOI:** 10.1101/2021.12.19.473380

**Authors:** Dhiraj Mannar, James W. Saville, Xing Zhu, Shanti S. Srivastava, Alison M. Berezuk, Katharine S. Tuttle, Citlali Marquez, Inna Sekirov, Sriram Subramaniam

## Abstract

The newly reported Omicron variant is poised to replace Delta as the most rapidly spread SARS-CoV-2 variant across the world. Cryo-EM structural analysis of the Omicron variant spike protein in complex with human ACE2 reveals new salt bridges and hydrogen bonds formed by mutated residues R493, S496 and R498 in the RBD with ACE2. These interactions appear to compensate for other Omicron mutations such as K417N known to reduce ACE2 binding affinity, explaining our finding of similar biochemical ACE2 binding affinities for Delta and Omicron variants. Neutralization assays show that pseudoviruses displaying the Omicron spike protein exhibit increased antibody evasion, with greater evasion observed in sera obtained from unvaccinated convalescent patients as compared to doubly vaccinated individuals (8-vs 3-fold). The retention of strong interactions at the ACE2 interface and the increase in antibody evasion are molecular factors that likely contribute to the increased transmissibility of the Omicron variant.

---

The Omicron (B.1.1.529.1) variant, first reported on November 24, 2021, was quickly identified as a variant of concern with potential to spread rapidly across the world. There is widespread concern about the speed with which the Omicron variant is currently circulating even amongst doubly vaccinated individuals. The Omicron variant spike protein has 3-5 times more mutations than that seen in any of the previous SARS-CoV-2 strains (*1*). Understanding the consequences of these mutations for ACE2 receptor binding and neutralizing antibody evasion is important in guiding the development of effective therapeutics to limit the spread of the Omicron and related variants.

The Omicron variant has 37 mutations (Figure 1A) in the spike protein relative to the initial Wuhan strain, with 15 of them present in the receptor binding domain (RBD) (*1*). The RBD is the region of the spike protein that mediates both attachment to human cells via the ACE2 receptor and is the primary target of neutralizing antibodies (*2, 3*). The Delta variant, which was the predominant SARS-CoV-2 lineage until the emergence of Omicron, has 7 mutations in the spike protein relative to the Wuhan strain but only two of these mutations (T478K and D614G) are in common with the Omicron strain. Analysis of the sequence of the Omicron genome suggests that it is not derived from any of the currently circulating variants, and may have a different origin (*4*).

**Figure 1.**
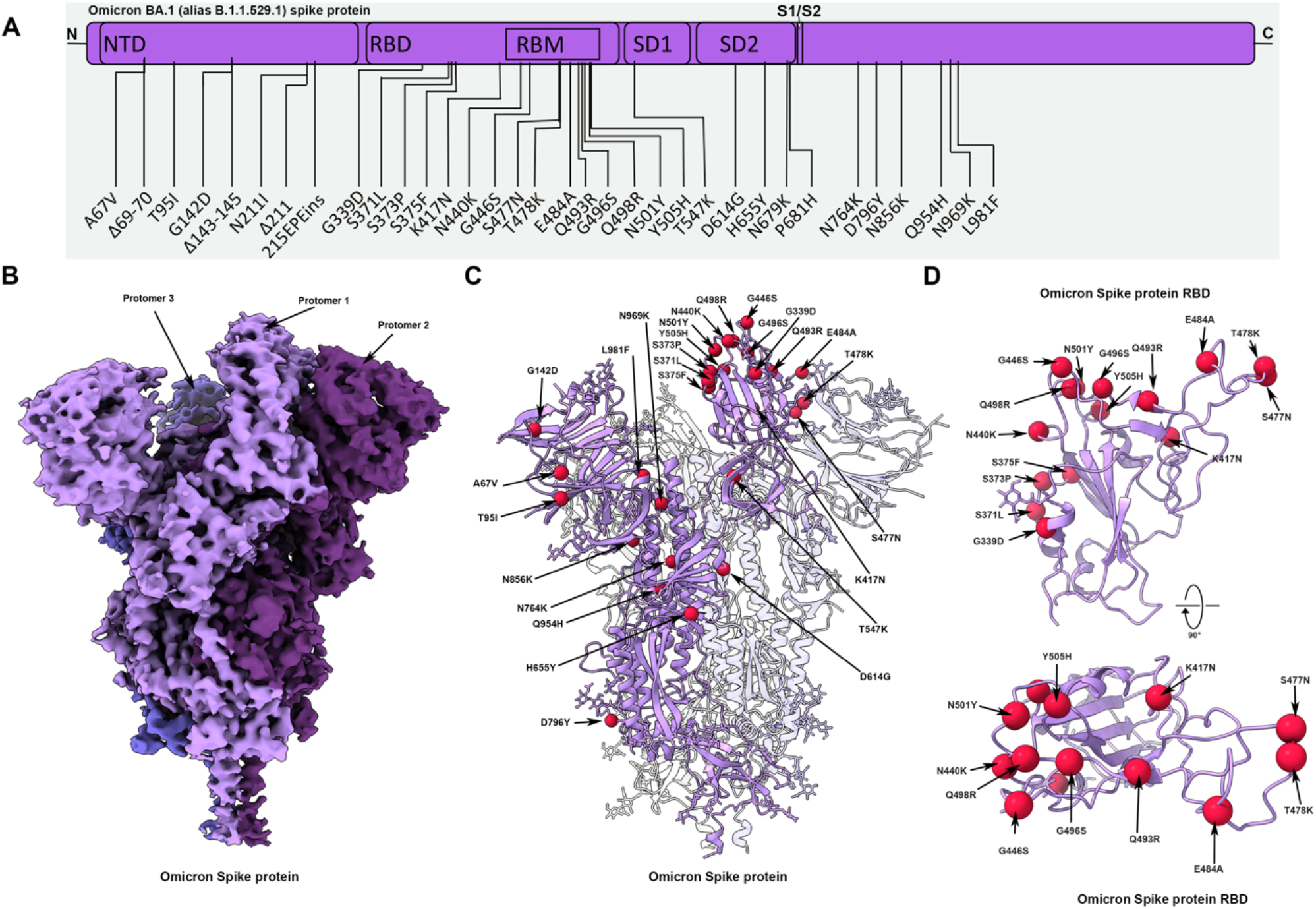
Cryo-EM structure of the Omicron spike protein. **(A)** A schematic diagram illustrating the domain arrangement of the spike protein. Mutations present in the Omicron variant spike protein are labeled. **(B)** Cryo-EM map of the Omicron spike protein at 2.79 Å. Protomers are coloured in shades of purple. **(C)** Cryo-EM structure of Omicron spike protein indicating the locations of all mutations on one protomer. **(D)** The Omicron spike receptor-binding domain (RBD) shown in two orthogonal orientations. Cα of all mutated residues are shown as red spheres.

Cryo-EM structural analysis of the Omicron spike protein ectodomain shows that the overall organization of the trimer is similar to that observed for the ancestral strain (*5-7*) and all earlier variants (*8-10*) (Figure 1B, Table S1). The RBD in one of the protomers (protomer 1) is well-resolved and is in the “down” position, while the other two RBDs are less well-resolved because they are flexible relative to the rest of the spike protein polypeptide. The mutations in the Omicron variant spike protein are distributed both on the surface and the interior of the spike protein (Figure 1C). Interestingly, the mutations in the RBD are predominantly distributed on one face of the domain (Figure 1D), which spans regions that bind ACE2 as well as those that form epitopes for numerous broadly neutralizing antibodies (*11*).

The Omicron variant shares RBD mutations in common with previous variants of concern (K417N, T478K, and N501Y). The N501Y and K417N mutations have been extensively individually characterized for their effect on ACE2 binding, and exhibit increased and decreased ACE2 binding affinity respectively (*8, 12-16*). These mutational effects have been demonstrated to act in a modular fashion, preserving the same general impact on ACE2 affinity when present in isolation or in combination with other RBD mutations (*12*). However, the Omicron RBD contains 12 additional mutations which have not yet been characterized in depth for their effect on ACE2 binding affinity, making it difficult to predict the overall change in ACE2 affinity for the Omicron variant. To measure the potential impact of Omicron mutations on the ACE2 binding affinity of its spike protein, we performed surface plasmon resonance studies and compared the resulting binding affinities (K_D_) to wild-type and Delta spikes (Figure 2). Wild-type (WT) is used in this work to refer to the ancestral Wuhan-1 strain with the addition of the D614G mutation. While the Omicron spike protein exhibits a measurable increase in affinity for ACE2 relative to the ancestral Wuhan strain (in agreement with a recent preprint (*17*)), the ACE2 affinity is similar for Delta and Omicron variants (Figure 2D). The Omicron variant includes the K417N mutation that is known to reduce ACE2 binding significantly (*12, 16*). The absence of a decrease in overall ACE2 binding affinity for the Omicron spike protein suggests there are likely compensatory mutations that restore higher affinity for ACE2, which should be possible to visualize in a high-resolution structure of the complex between the spike protein and ACE2.

**Figure 2.**
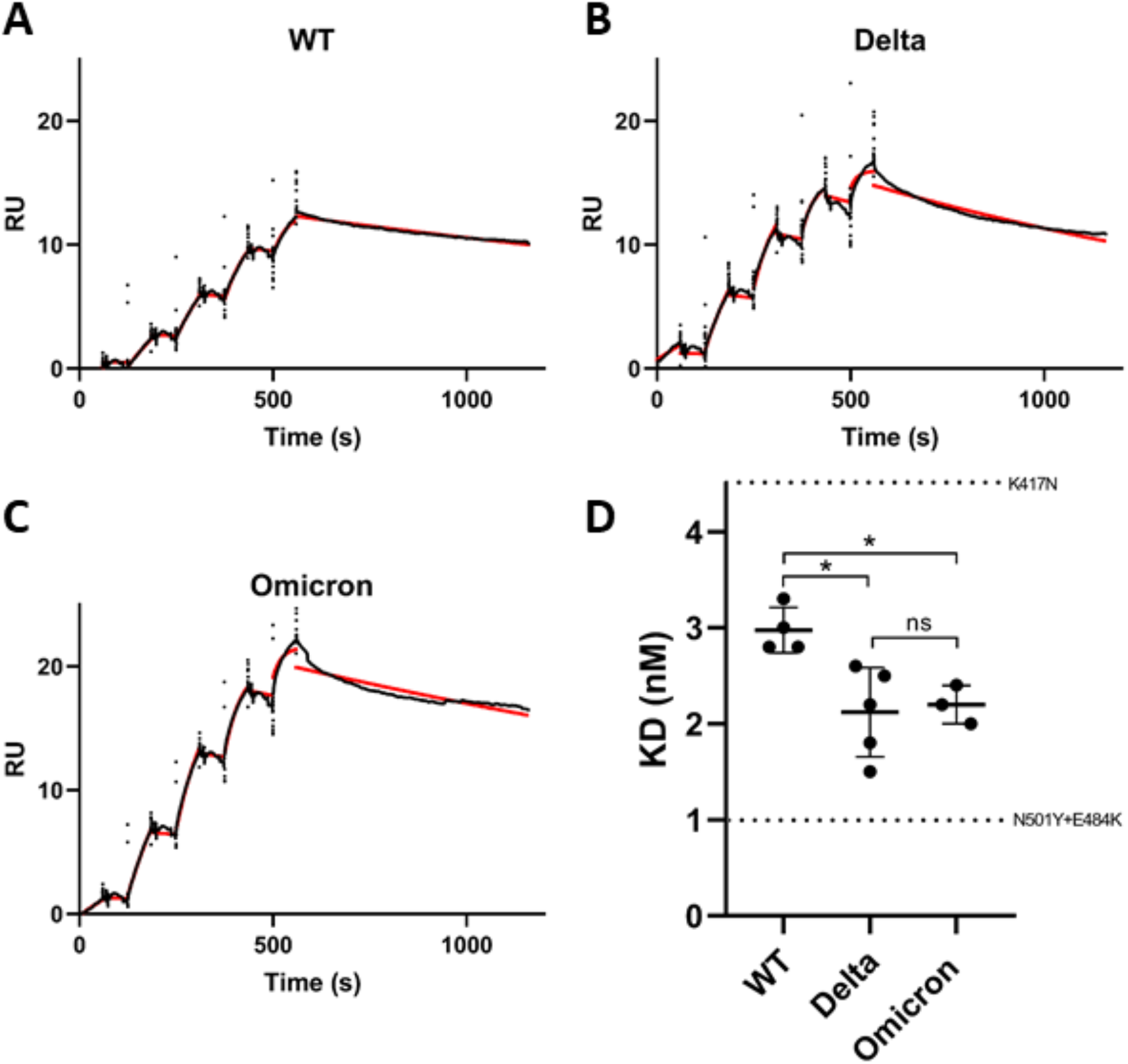
SPR analysis of the wild-type, Delta, and Omicron spike protein affinities for ACE2. **(A-C)** Representative curves of single-cycle kinetic analyses of S protein – ACE2 binding. Red curves show the fitting to the raw data (shown in black) using a 1:1 binding model. 6.25, 31.25, 62.5, 125, 250 nM of each spike protein was injected in successive cycles. (RU: Response units). **(D)** Quantitation of dissociation constants (KD) for the wild-type, Delta, and Omicron S protein – ACE2 interaction. The standard deviation of at least three replicates is shown. Horizontal dotted lines are plotted for the K417N and N501Y + E484K individually mutated S proteins to demonstrate the range of this assay (See figure S2 for binding data). A Tukey’s multiple comparisons test was performed on the wild-type, Delta, and Omicron binding affinities (*P≤0.05, ns = not significant).

Cryo-EM structural analysis of the ACE2-Omicron spike protein complex shows strong density for ACE2 bound to the RBD of one of the protomers in the “up” position (Figure 3A, Table S1). Weaker density is observed for a second bound ACE2, suggesting partial occupancy of the second RBD under the conditions of our experiment. The stoichiometry of ACE2 binding can be variable depending on the experimental conditions; in this analysis we have concentrated on the structure of the ACE2-spike protein interface in the most strongly bound ACE2 molecule. Focused refinement of the RBD-ACE2 region resulted in a density map with a resolution of 2.66 Å at the spike protein-ACE2 interface (Figure 3B), allowing visualization of sidechains involved in the interface (Figure 3C). In Figure 3D, we compare the key interactions at this interface in the Omicron variant with corresponding interactions that we have reported recently for the Delta variant ((*18*), accepted at Nature Communications). In the Delta variant-ACE2 complex, there are hydrogen bonds formed by residues Q493 and Q498 on the spike protein with residues E35 and Q42, respectively, on ACE2. In the Omicron variant, three mutations are observed in this stretch: Q493R, G396S and Q498R. Residue R493 replaces the hydrogen bond to ACE2 residue E35 with a new salt bridge, while residue R498 forms a new salt bridge with ACE2 residue D38 in addition to maintaining a hydrogen bonded interaction between residue 498 and ACE2 residue Q42. RBD residue S496 adds a new interaction at the interface by forming a hydrogen bond with ACE2 residue K353.

**Figure 3.**
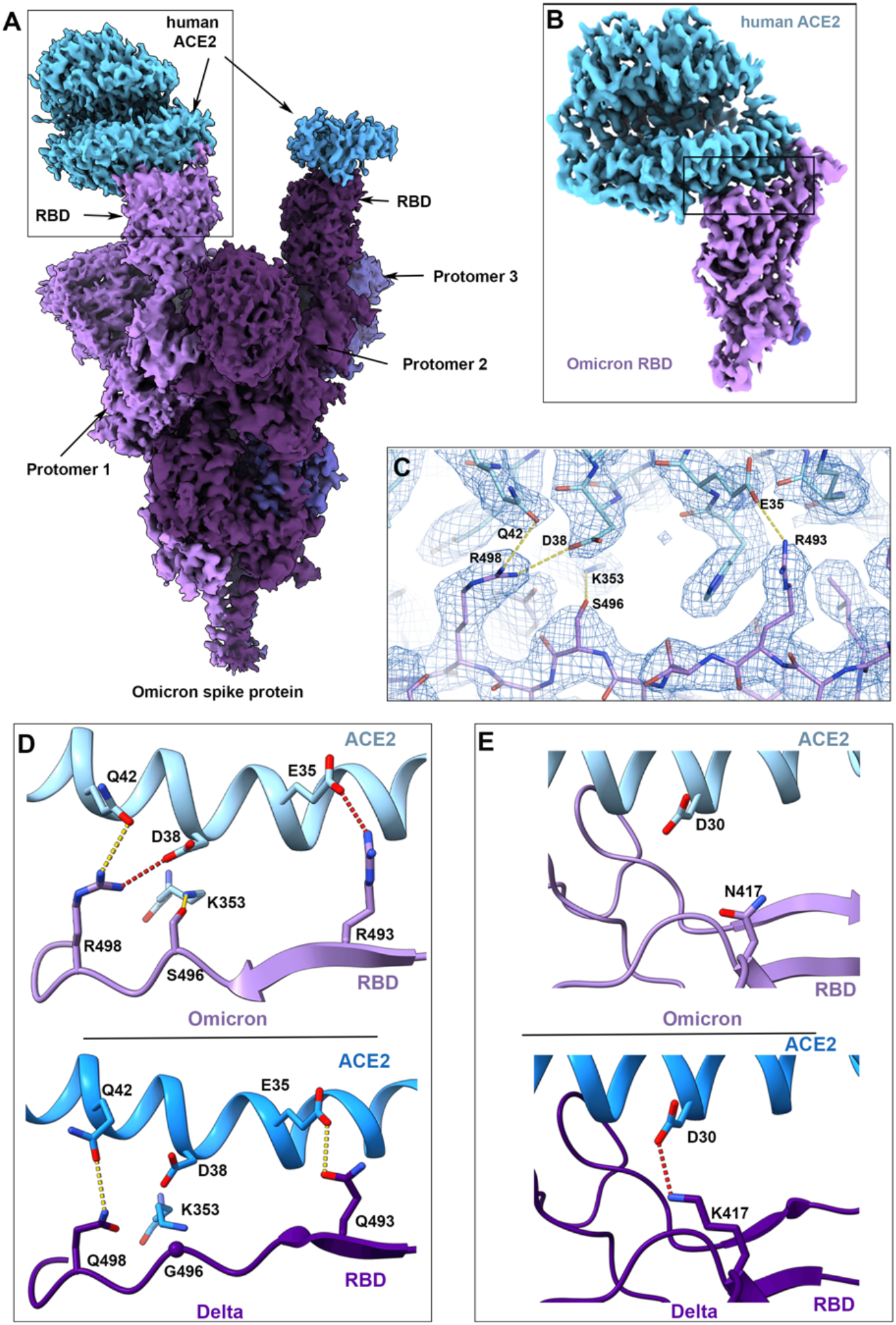
Cryo-EM structure of the Omicron spike protein-ACE2 complex. **(A)** Cryo-EM map of the Omicron spike protein in complex with human ACE2 at 2.45 Å resolution after global refinement. The three protomers are colored in shades of purple and the density for bound ACE2 proteins is coloured in blue. **(B)** Cryo-EM map of the Omicron spike RBD in complex with ACE2 at 2.66 Å resolution after focused refinement. The inset box indicates the region highlighted in (C). **(C)** Cryo-EM density at the Omicron spike RBD-ACE2 interface, with fitted atomic model. Yellow dashed lines represent hydrogen bonds. **(D-E)** Comparison of the RBD-ACE2 interface between the Omicron (upper) and Delta (lower) variants. Compared to the Delta variant, new interactions are formed as a result of the mutations Q493R, G496S and Q498R (D), and the salt bridge between RBD K417 and ACE2 D30 is lost (E) in the Omicron variant. Yellow and red dashed lines represent hydrogen bonds and ionic interactions respectively.

While these new interactions formed by residues 493, 496 and 498 likely make spike-ACE2 binding stronger, this is offset by the loss of a key salt bridge between spike protein residue K417 and ACE2 residue D30 that is present in the Delta variant. In isolation, the K417N mutant displays reduced ACE2 binding affinity (*12, 16*), but our findings suggest that the new mutations in the Omicron interface have a compensatory effect on the strength of ACE2 binding, providing an explanation for the similar ACE2 binding affinities observed (Figure 2).

We next investigated the effects of Omicron mutations on neutralization by (i) a selection of monoclonal antibodies, (ii) sera obtained from 30 doubly vaccinated individuals with no prior history of COVID-19 infection and (iii) sera obtained from a set of 68 unvaccinated convalescent patients who recovered from infection by either Alpha, Gamma or Delta variants (summary of patient demographics is in Table S2). We performed neutralization experiments using pseudovirions that incorporate the wild-type, Delta, or Omicron variant S proteins and compared the resulting antibody evasion of viral entry. The comparison to evasion relative to the Delta variant is likely the most important and relevant given that the Omicron variant is rapidly supplanting the Delta variant in global prevalence, though comparison to wild-type SARS-CoV-2 is still relevant given the majority of current SARS-CoV-2 vaccine immunogens are based on this sequence(*19*).

We used a panel of neutralizing monoclonal antibodies that include RBD-directed antibodies (ab1, ab8, S309, S2M11;(*20-23*)) and two N-terminal domain (NTD)-directed antibodies (4-8 and 4A8; (*24, 25*)) to investigate the impact of Omicron RBD and NTD mutations on monoclonal antibody escape. In contrast to the Alpha, Beta, Gamma, Kappa, Epsilon, and Delta variants of SARS-CoV-2, the Omicron variant exhibited measurable evasion from every antibody in this panel with complete escape from five of the six antibodies tested (Figure 4A) (*26*). The neutralizing activity of both the NTD-directed antibodies (4-8 and 4A8) is completely knocked out for the Omicron variant, as seen previously in the Alpha variant which contains identical or similar deletions in its NTD (Δ69-70 and Δ144-145). The Omicron variant does not show complete escape from S309, an antibody undergoing evaluation in clinical trials to treat patients with COVID-19 although there is still a 4-fold decrease in its neutralization potency relative to the ancestral strain(*27*). The unusually high number of mutations in the Omicron variant spike protein thus appear to confer unprecedently broad antibody escape relative to previously emerged variants of SARS-CoV-2 (*17*).

**Figure 4.**
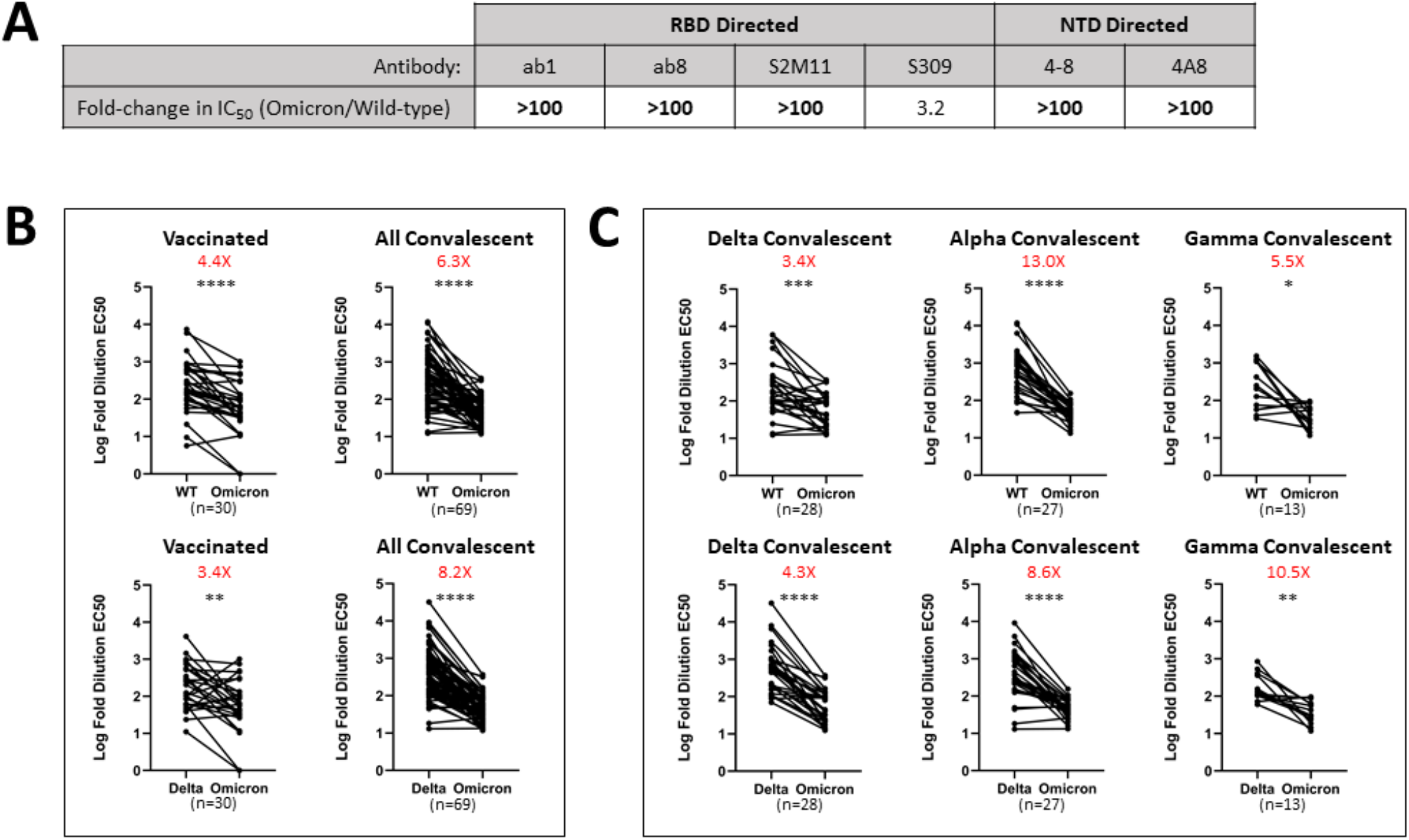
Monoclonal antibodies and vaccinated and convalescent patient-derived sera exhibit decreased Omicron neutralization potency. **(A)** Fold-changes in monoclonal antibody neutralization for Omicron variant pseudoviruses relative to wild-type (D614G). **(B)** Log-fold EC50 dilutions for vaccinated and convalescent patient sera for either wild-type (D614G) versus Omicron variant pseudoviruses (top) or Delta and Omicron variant pseudoviruses (bottom). **(C)** As in **(B)** with a breakdown of the convalescent patients into previous infection with Delta, Alpha, and Gamma variants of concern. A pairwise statistical significance test was performed using the Wilcoxon matched pairs test (*P≤0.05, **P≤0.01, ***P≤0.001, ****P≤0.0001). Each data point is the average of n = 2 replicates. The fold-change in the geometric mean between the two groups is shown in red text at the top of each plot.

Sera from convalescent patients displayed on average a 6.3x decrease in ability to neutralize the Omicron variant relative to the wild-type (Figure 4B upper panel). Sera from the vaccinated cohort also displayed reduced neutralization ability (4.4x decrease) with a wider variation driven by some individuals that showed exceptional loss of neutralization ability to Omicron. The comparison of change in neutralization potential between Delta and Omicron variants is a more relevant comparison given the world-wide dominance of Delta. Sera from convalescent patients shows an even greater drop in neutralization potency relative to the Delta variant (8.2x decrease) while the vaccinated group also shows reduction in potency, although to a lesser extent (3.4x decrease). We note that the majority of the doubly vaccinated cohort consisted of individuals who were vaccinated with a schedule of at least 8 weeks between doses, recently shown to generate a better humoral response than the manufacturer-recommended interval of 3-4 weeks (*28*). The longer interval between doses could result in less pronounced reduction in neutralization of the Omicron variant.

A finer analysis of the unvaccinated convalescent cohort stratified into those who recovered from infection by either Delta, Alpha, or Gamma variants (figure 4C) reinforces the reduction in neutralization potency against the Omicron variant in all populations, with especially striking drops for patients who recovered from infection from the earlier Alpha and Gamma variants. The findings we report here are consistent with several other recent reports (*17, 29-31*) that the Omicron variant is more resistant to neutralization than any other variant of concern that has emerged over the course of the pandemic.

The mechanisms underlying the rapid spread of the newly emerged Omicron variant are of fundamental interest given the likelihood that this virus could become the dominant variant of SARS-CoV-2. The large number of mutations on the surface of the spike protein including the immunodominant RBD (Figure 1) would be expected to help the virus evade from antibodies elicited by vaccination or prior infection. However, it is remarkable that the Omicron variant evolved to retain its ability to bind ACE2 efficiently despite these extensive mutations. The cryo-EM structure of the spike protein-ACE2 complex provides a structural rationale for how this is achieved: interactions involving the new mutations in the Omicron variant at residues 493, 496 and 498 appear to restore ACE2 binding efficiency that would be lost due to other mutations such as K417N. The Omicron variant thus appears to have evolved to selectively balance two critical features, namely the increase in escape from neutralization but without compromising its ability to interact efficiently with ACE2. The increase in antibody evasion and the retention of strong interactions at the ACE2 interface are thus likely to represent key molecular features that contribute to the increase in transmissibility of the Omicron variant.

## DATA AVAILABILITY

Cryo-EM reconstructions and atomic models generated during this study are available at the PDB and EMBD databases under the following accession codes: Unbound Omicron spike protein trimer (PDB ID 7T9J, EMD-EMDB 25759), global ACE2-bound Omicron spike protein trimer (PDB ID 7T9K, EMD-25760), focus-refinement of the ACE2-RBD interface for the ACE2-bound Omicron spike protein trimer (PDB ID 7T9L, EMD-25761).

## ACKNOWLEDGMENTS

This work was supported by awards to S.S. from a Canada Excellence Research Chair Award, the VGH Foundation, Genome BC, Canada, and from the Tai Hung Fai Charitable Foundation. D.M. is supported by a CIHR Frederick Banting and Charles Best Canada Graduate Scholarship Master’s Award (CGS-M). J.W.S is supported by a CIHR Frederick Banting and Charles Best Canada Graduate Scholarships Doctoral Award (CGS D) and a UBC President’s Academic Excellence Initiative PhD Award. We thank Dr. Karoline Leopold (UBC) and Dr. Charles Leung (Gandeeva Therapeutics Inc.) for assistance with the surface plasmon resonance experiments and for helpful discussions.

## AUTHOR CONTRIBUTIONS

This work was the result of a concerted team effort from all individuals listed as authors. J.W.S. and D.M., carried out expression and purification of the Omicron spike protein and antibodies. D.M. performed the SPR binding analyses. D.M. and J.W.S. performed the pseudovirus neutralization experiments. I.S. and C.M. provided the vaccine-induced patient-derived sera samples and aided the interpretation of the patient data. A.M.B., S.S.S. and K.S.T. carried out experimental aspects of electron microscopy including specimen preparation and data collection. X.Z. carried out computational aspects of image processing and structure determination. X.Z., S.S.S., D.M., J.W.S., and S.S. interpreted and analyzed the cryo-EM structures, binding analyses and patient neutralization data and composed the manuscript with input from the rest of the authors. S.S. provided overall supervision for the project.

## Method details

### Ethics Statement

Patient derived sera samples were collected according to the CARE COVID Study (http://www.bccdc.ca/health-professionals/clinical-resources/covid-19-care/covid-19-serology-care-covid-study) with ethics approval from the UBC Clinical Research Ethics Board.

### Expression and Purification of Omicron Recombinant Spike Protein Constructs

The production of the SARS-CoV-2 wild-type (D614G), K417N, N501Y+E484K and Delta Hexapro S proteins were described previously (*12,17*).

The SARS-CoV-2 Omicron HexaPro S protein gene was synthesized and inserted into pcDNA3.1 (GeneArt Gene Synthesis, Thermo Fisher Scientific).

Expi293F cells (Thermo Fisher, Cat# A14527) were grown in suspension culture using Expi293 Expression Medium (Thermo Fisher, Cat# A1435102) at 37**°**C, 8% CO_2_. Cells were transiently transfected at a density of 3 × 10^6 cells/mL using linear polyethylenimine (Polysciences Cat# 23966-1). The media was supplemented 24 hours after transfection with 2.2 mM valproic acid, and expression was carried out for 3 days at 37**°**C, 8% CO_2_. The supernatant was harvested by centrifugation and filtered through a 0.22-μM filter prior to loading onto a 5 mL HisTrap excel column (Cytiva). The column was washed for 20 CVs with wash buffer (20 mM Tris pH 8.0, 500 mM NaCl), 5 CVs of wash buffer supplemented with 20 mM imidazole, and the protein eluted with elution buffer (20 mM Tris pH 8.0, 500 mM NaCl, 500 mM imidazole). Elution fractions containing the protein were pooled and concentrated (Amicon Ultra 100 kDa cut off, Millipore Sigma) for gel filtration. Gel filtration was conducted using a Superose 6 10/300 GL column (Cytiva) pre-equilibrated with GF buffer (20 mM Tris pH 8.0, 150 mM NaCl). Peak fractions corresponding to soluble protein were pooled and concentrated to 4.5–6.5 mg/mL (Amicon Ultra 100 kDa cut off, Millipore Sigma). Protein samples were flash-frozen in liquid nitrogen and stored at −80**°**C.

### Antibody Production

VH-FC ab8, IgG ab1, Fab S309, Fab S2M11, Fab 4-8, and Fab 4A8 were produced as previously described (*19-20*).

### Surface Plasmon Resonance

SPR experiments were performed on the Biacore T200 instrument. Recombinant mouse ACE2-mFc (SinoBiological) was immobilized using the series S protein A chip. Increasing concentrations (6.25nM, 31.25nM, 62.5nM, 125nM, 250nM) of various spike protein trimers were flowed over the surface for single cycle kinetic experiments. The surface was regenerated in 10mM glycine pH 1. The experiments were performed at 25 degrees Celsius, using 10mM HEPES, 150mM NaCl, 3mM EDTA and 0.05% v/v Surfactant P20 as running buffer. Reference-subtracted curves were fitted to a 1:1 binding model using Biacore evaluation software.

### Pseudovirus Neutralization Assay

SARS-CoV-2 S protein Omicron genes were synthesized and inserted into pcDNA3.1 (GeneArt Gene Synthesis, Thermo Fisher Scientific). Pseudotyped retroviral particles were produced in HEK293T cells as described previously (*32*). Briefly, a lentiviral packaging system was utilized in combination with plasmids encoding the full-length SARS-CoV-2 wild-type (D614G), Delta, and Omicron spikes, along with a transfer plasmid encoding luciferase and GFP as a dual reporter gene. Pseudoviruses were harvested 60 h after transfection and filtered with a 0.45 µm PES filter. For neutralization assays, HEK293T-ACE2-TMPRSS2 cells (*33*) (BEI Resources cat# NR-55293) were seeded in 384-well plates at 20 000 cells. The next day, pseudovirus preparations normalized for viral capsid p24 levels (Lenti-X™ GoStix™ Plus) were incubated with dilutions of the indicated antibodies, sera, or media alone for 1 h at 37°C prior to addition to cells and incubation for 48 h. Cells were then lysed and luciferase activity assessed using the ONE-Glo™ EX Luciferase Assay System (Promega) according to the manufacturer’s specifications. Detection of relative luciferase units was carried out using a Varioskan Lux plate reader (Thermo Fisher).

### Electron Microscopy Sample Preparation and Data Collection

For cryo-EM, 2.25 mg/mL S protein and S protein-ACE2 complex (1:2.3 S protein trimer:ACE2 molar ratio) samples were vitrified using a Vitrobot Mark IV (Thermo Fisher Scientific) plunge freezing device. Quantifoil R1.2/1.3 Cu mesh 200 holey carbon grids were first glow discharged for 20 seconds using a Pelco easiGlow glow discharge unit (Ted Pella) and then 1.8 µL of protein suspension was applied to the surface of the grid at a temperature of 10°C and a humidity level of >98%. Grids were then blotted (12 sec, blot force −10) and plunge frozen into liquid ethane. S protein-ACE2 complex grids were imaged using a 300 kV Titan Krios G4 transmission electron microscope (Thermo Fisher Scientific) equipped with a Falcon4 direct electron detector in electron event registration (EER) mode. Movies were collected at 155,000x magnification (calibrated pixel size of 0.5 Å per physical pixel) over a defocus range of −0.5 µm to −2 µm with a total dose of 40 e^-^/Å^2^ using EPU automated acquisition software. Grids containing the Omicron S protein alone were imaged using a 200 kV Glacios transmission electron microscope (Thermo Fisher Scientific) equipped with a Falcon4 camera operated in EER mode. Micrographs were collected using EPU at 190,000x magnification (physical pixel size 0.7 Å) over a defocus range of −0.5 µm to −2 µm with a total accumulated dose of 40 e^-^/Å^2^.

### Image Processing

The detailed data processing workflow is summarized in Supplementary Figures S1,S3. All data processing was done in cryoSPARC v.3.3.1 (*34*). Motion correction in patch mode (EER upsampling factor 1, EER number of fractions 40), CTF estimation in patch mode, blob particle picking, and particle extraction (box size 400 Å) were performed on-the-fly in cryoSPARC. Then particles were subjected to multiple rounds of 3D heterogeneous classification. The final 3D refinement was performed with per particle CTF estimation and aberration correction. For the complexes of Omicron spike protein ectodomain and human ACE2, local refinement was performed with a soft mask covering a single RBD and its bound ACE2.

### Model Building and Refinement

For models of Omicron spike protein ectodomain alone, the SARS-CoV-2 HexaPro S trimer with N501Y mutation (PDB code 7MJG) was fitted into the map using UCSF Chimera v.1.15 (*35*). Then, mutation and manual adjustment were carried out with COOT v.0.9.3 (*36*), followed by iterative rounds of real-space refinement in COOT and Phenix v.1.19 (*37*). Glycans were added at N-linked glycosylation sites in COOT. For models of Omicron spike-ACE2 complex, the RBD-ACE2 subcomplex was built using published coordinates (PDB code 7MJN) as the initial model, followed by refinement against local refinement maps. The obtained model was then docked back into global refinement maps together with the other individual domains of the spike protein. Model validation was performed using MolProbity (*38*). Figures were prepared using UCSF Chimera, UCSF ChimeraX v.1.1.1 (*39*), and PyMOL (v.2.2 Schrodinger, LLC).

**Figure S1.**
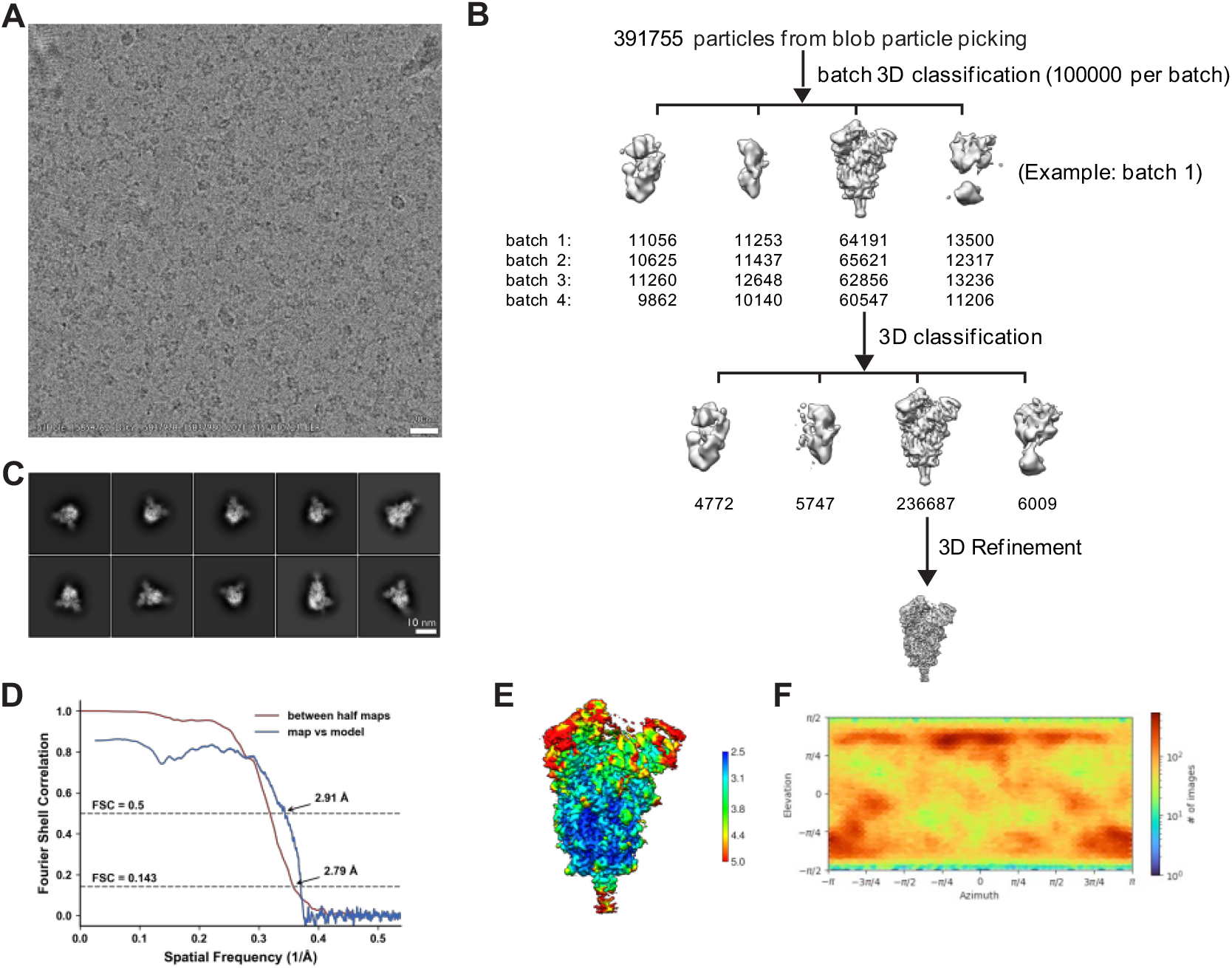
Cryo-EM data processing and validation for the Omicron spike protein ectodomain. (A) Representative cryo-EM micrograph. (B) Workflow of cryo-EM image processing. (C) Representative 2D classes. (D) FSC curves. (E) Local resolution. (F) Viewing direction distribution plot.

**Figure S2.**
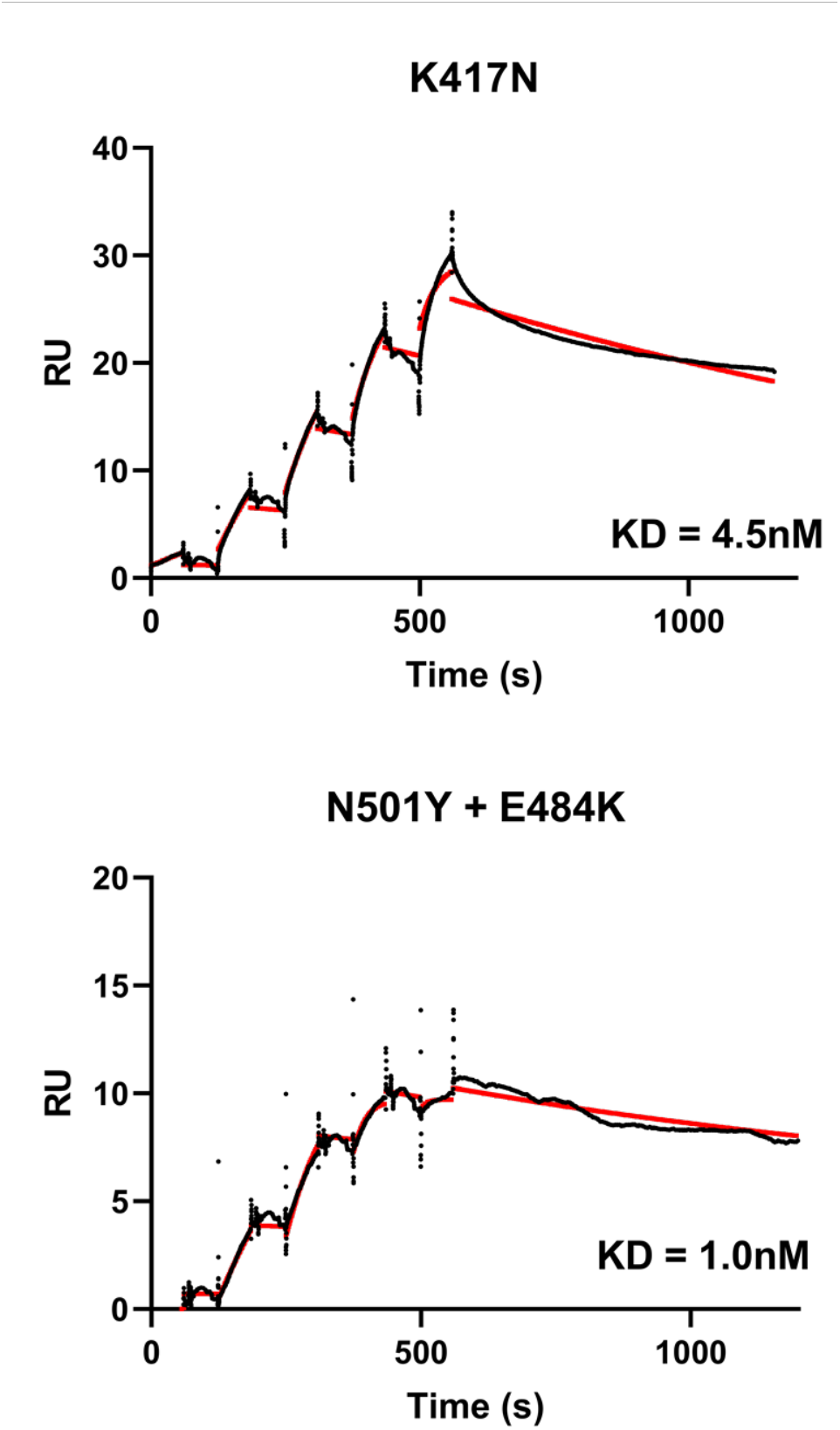
SPR Analysis of the K417N and N501Y+E484K spike protein affinities for ACE2. Red curves show the fitting to the raw data (shown in black) using a 1:1 binding model. 6.25, 31.25, 62.5, 125, 250 nM of each spike protein was injected in successive cycles. (RU: Response units).

**Figure S3.**
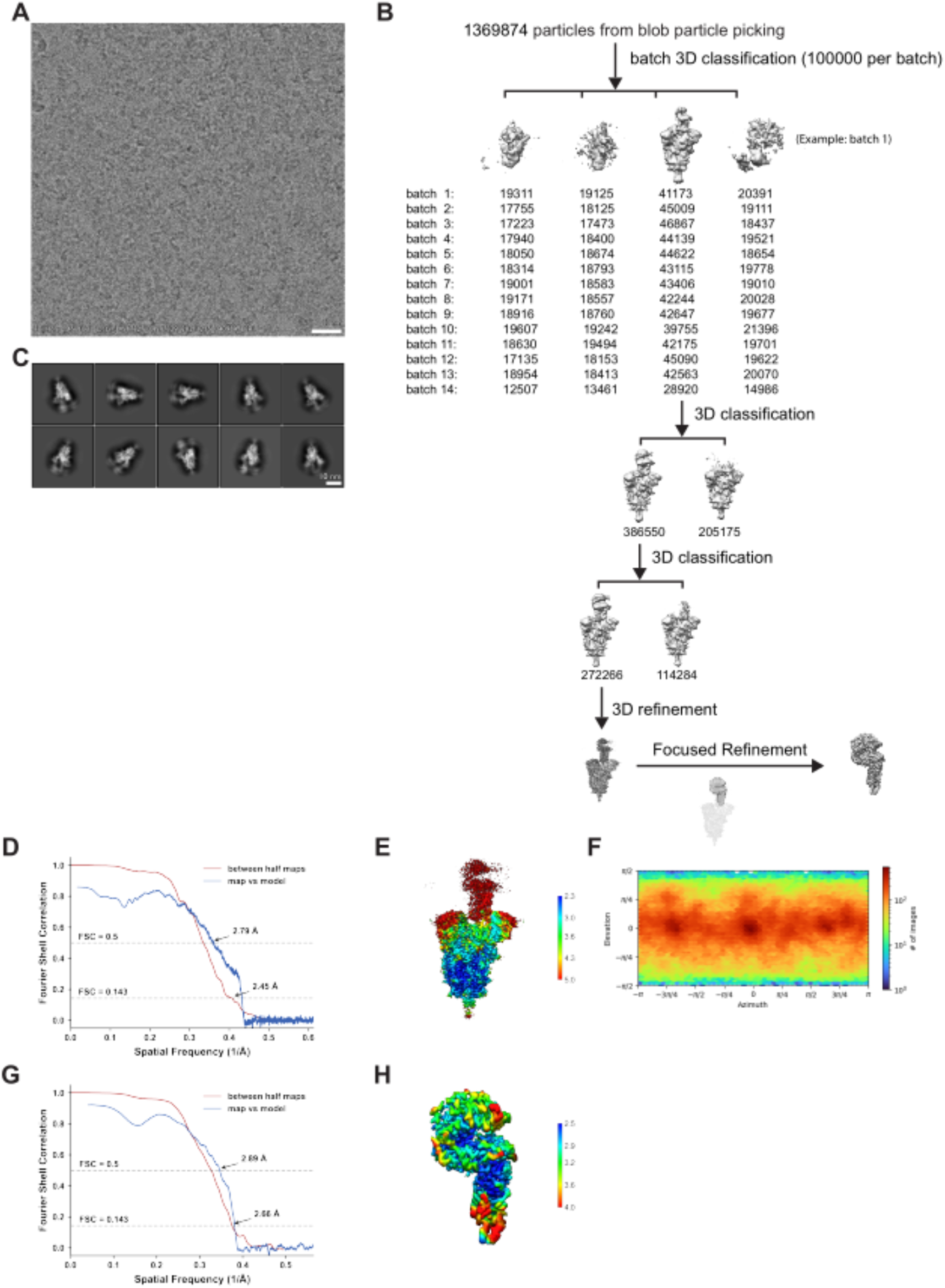
Cryo-EM data processing and validation for complex of Omicron spike protein ectodomain and human ACE2. (A) Representative cryo-EM micrograph. (B) Workflow of cryo-EM image processing. (C) Representative 2D classes. (D-F) FSC curves (D), local resolution (E) and viewing direction distribution plot (F) of global refinement. (G-H) FSC curves (G) and local resolution (H) of focused refinement.

**Figure S4.**
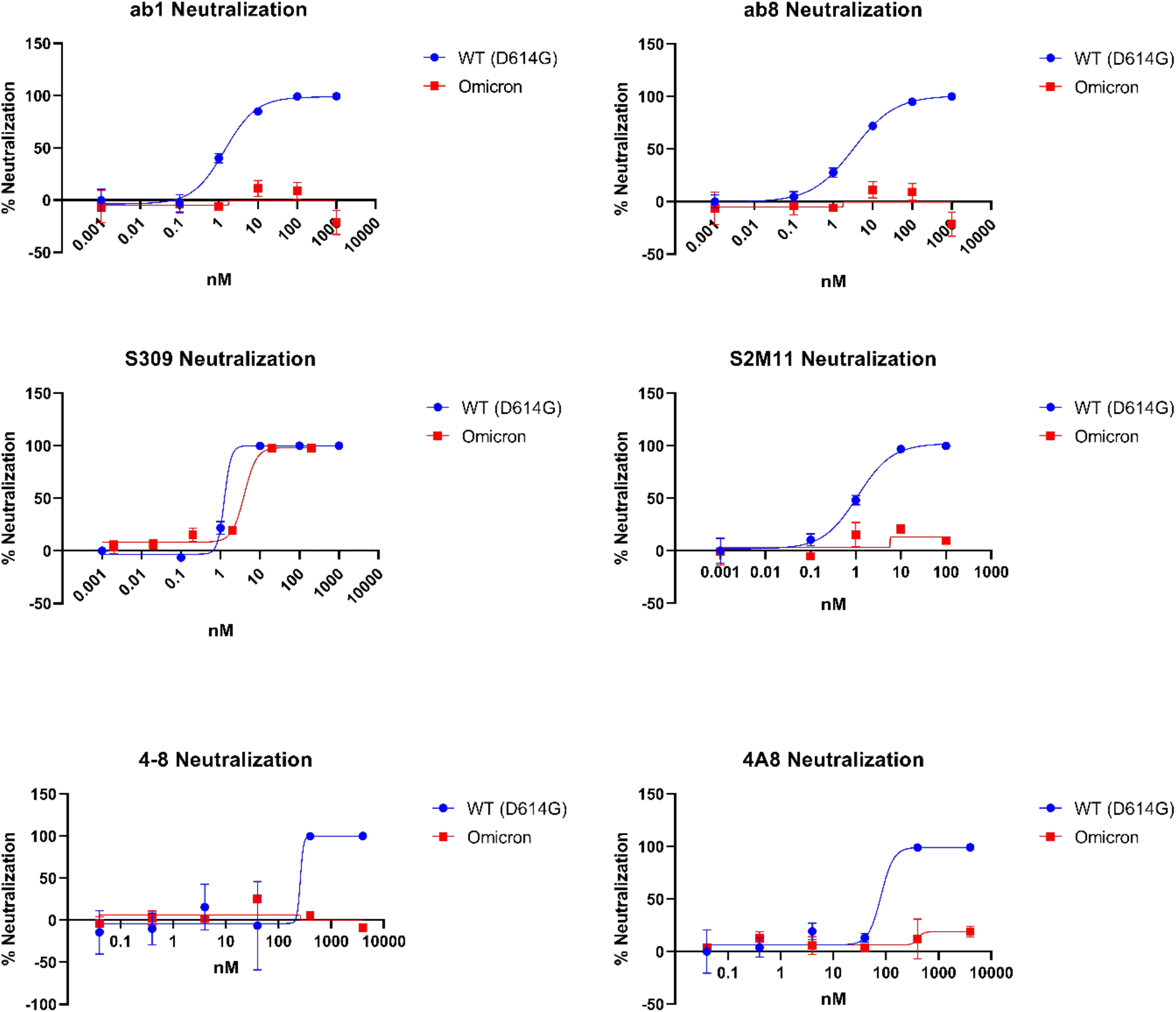
Monoclonal antibody neutralization of Omicron S protein pseudotyped viruses and comparison to previously determined^1^ wild-type (D614G) S protein pseudotyped virus neutralization curves. Points denote the mean of (n = 3) replicates, error bars denote the standard error of the mean

**Figure S5.**
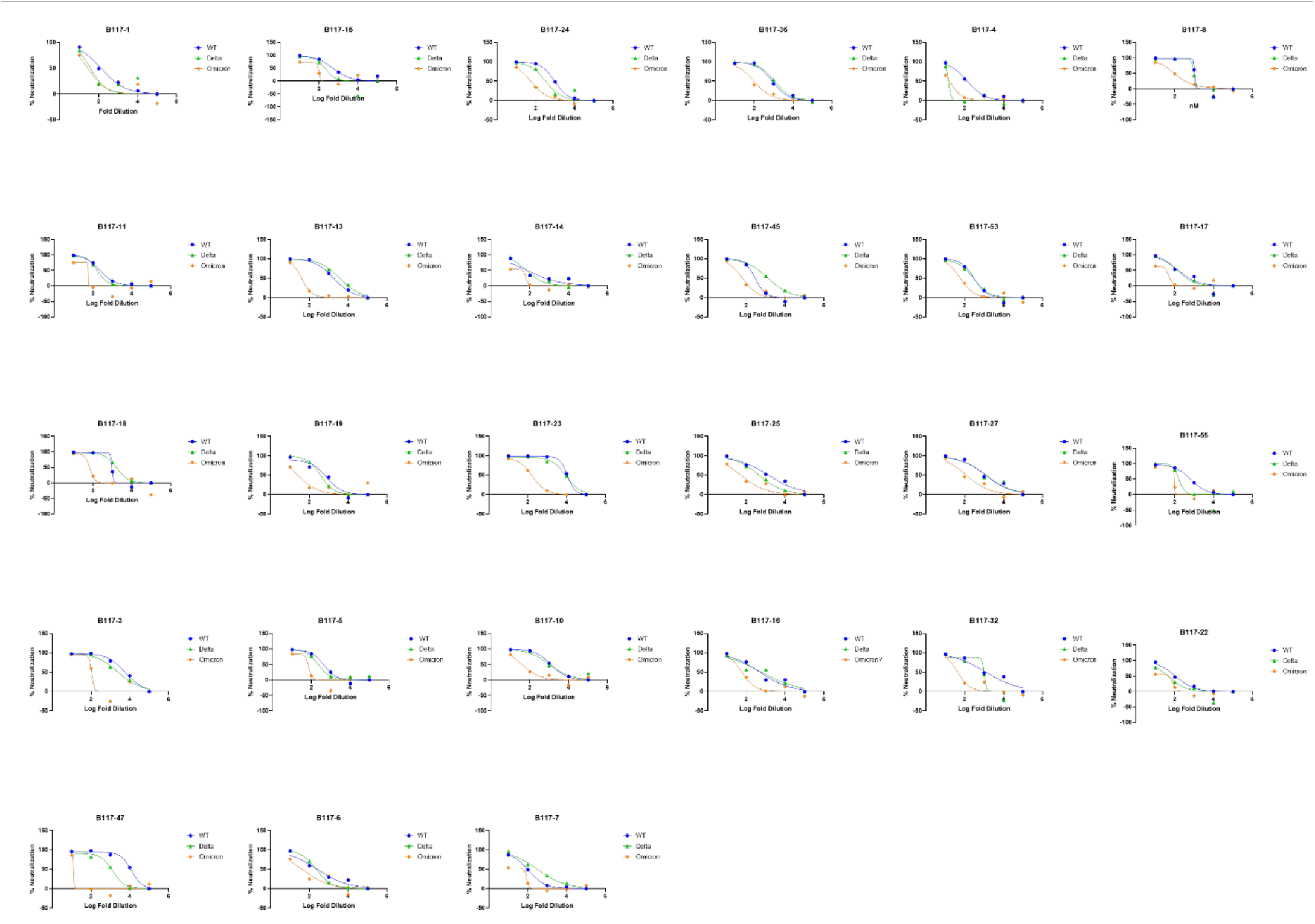
Data for the neutralization of Wild-type (D614G), Delta and Omicron S protein pseudotyped viruses with convalescent sera samples from patients previously infected with the Alpha variant. Data show the average of (n=2) replicates.

**Figure S6.**
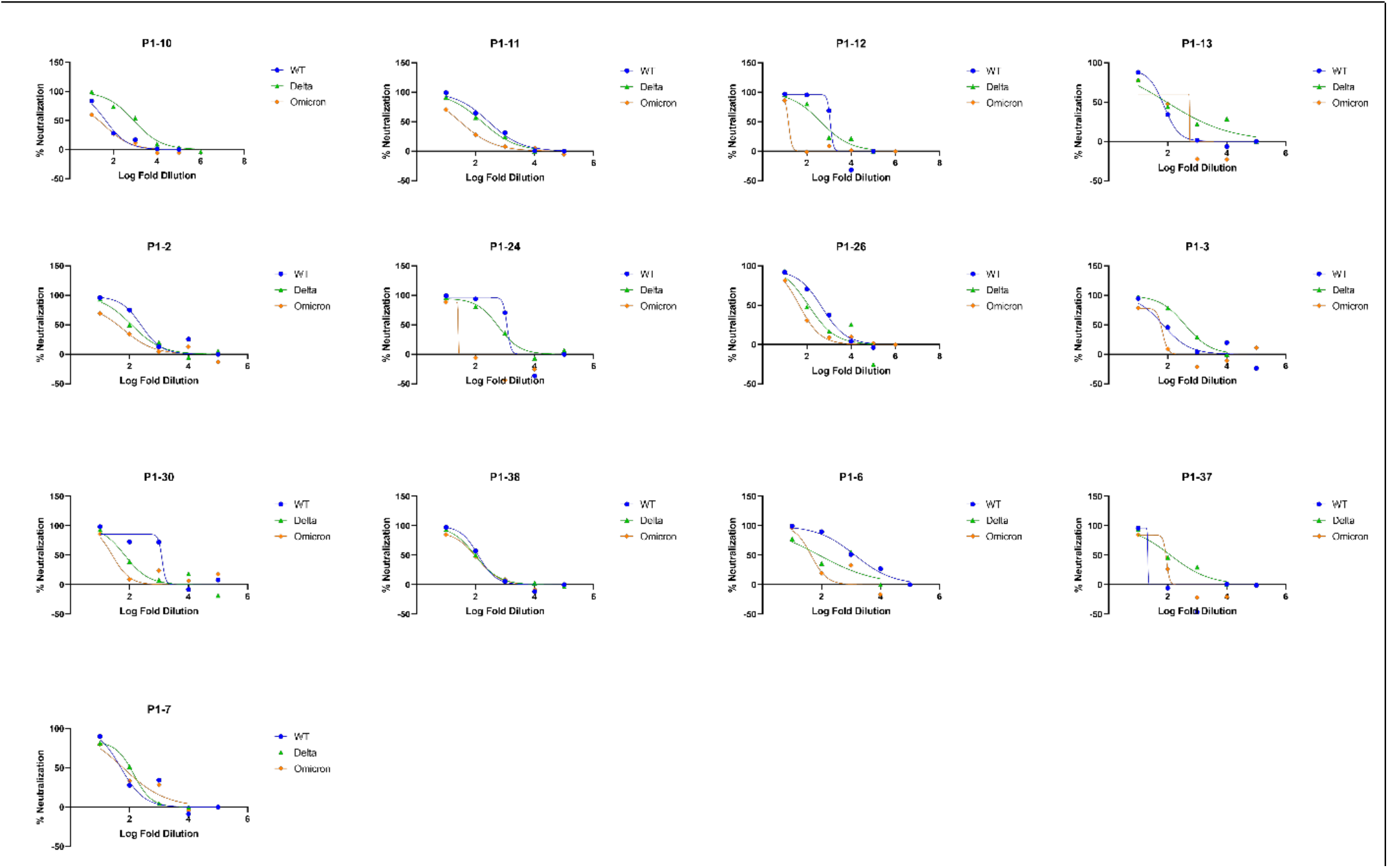
Data for the neutralization of Wild-type (D614G), Delta and Omicron S protein pseudotyped viruses with convalescent sera samples from patients previously infected with the Gamma variant. Data show the average of (n=2) replicates.

**Figure S7.**
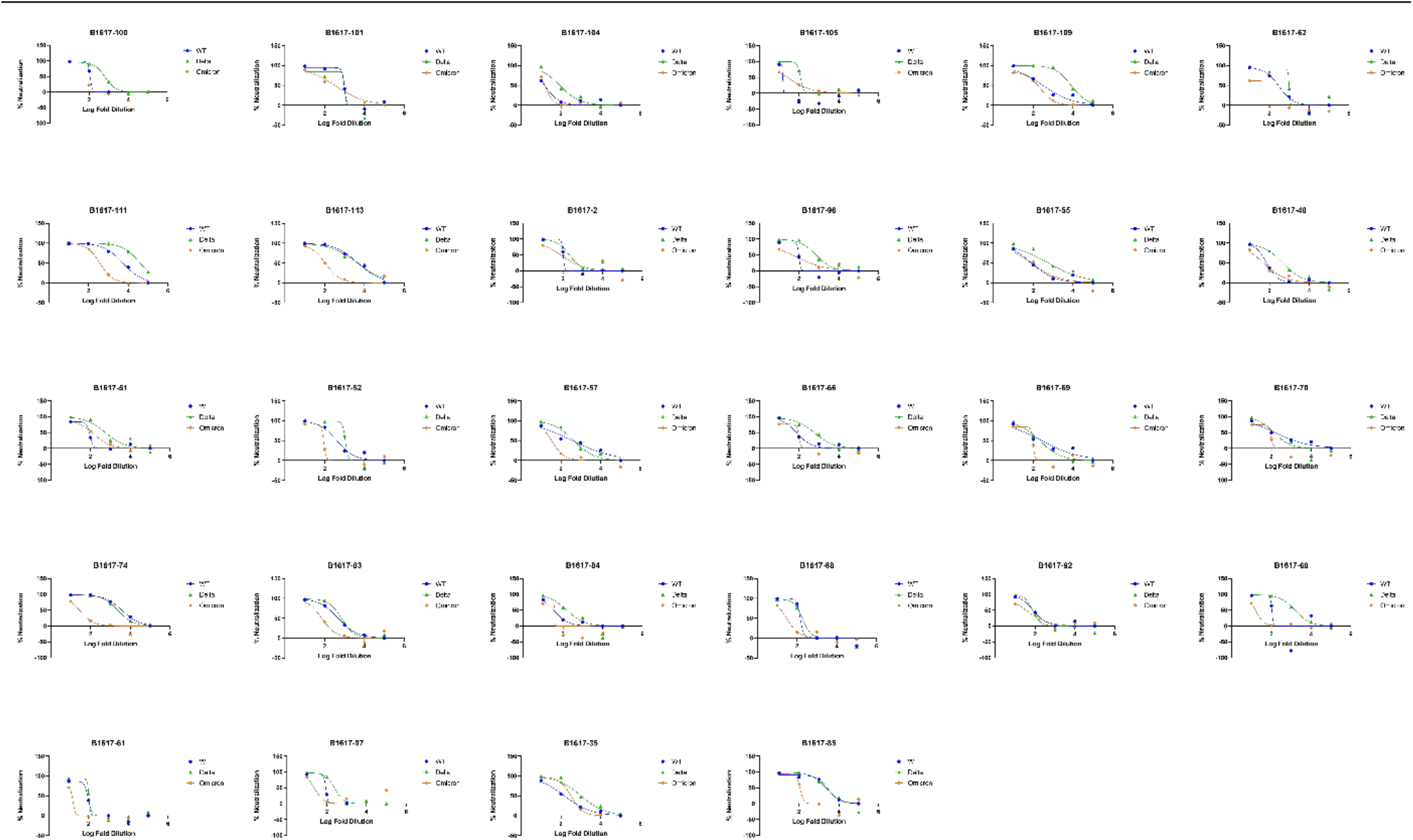
Data for the neutralization of Wild-type (D614G), Delta and Omicron S protein pseudotyped viruses with convalescent sera samples from patients previously infected with the Delta variant. Data show the average of (n=2) replicates.

**Figure S8.**
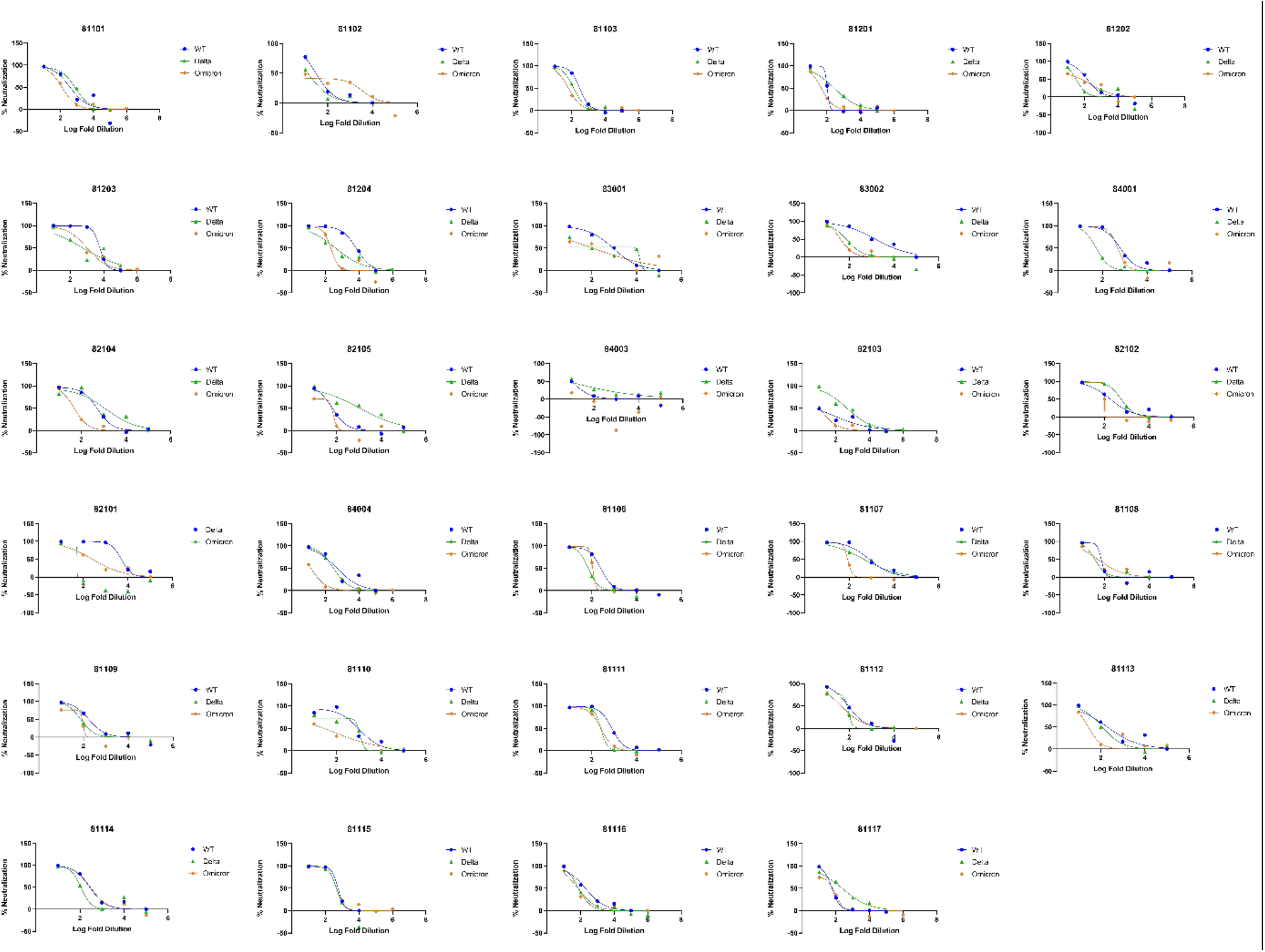
Data for the neutralization of Wild-type (D614G), Delta and Omicron S protein pseudotyped viruses with convalescent sera samples from non convalescent patients who received 2 vaccine doses. Data show the average of (n=2) replicates.

**Table S1:** CryoEM data collection, processing, refinement, and validation parameters for the structures reported in this publication.

|  | S(Omicron) | S(Omicron) + ACE2 |  |
| --- | --- | --- | --- |
|  | (EMDB 25759) | global refinement | focus refinement |
|  | (PDB 7T9J) | (EMDB 25760) | (EMDB 25761) |
|  |  | (PDB 7T9K) | (PDB 7T9L) |
| Data collection |  |  |  |
| Microscope | Glacios | Titan Krios G4 |  |
| Detector | Falcon4 | Falcon4 |  |
| Voltage (kV) | 200 | 300 |  |
| Nominal magnification | 190,000 | 155,000 |  |
| Defocus range (μm) | -2.0 to -0.5 | -2.0 to -0.5 |  |
| Physical pixel (Å) | 0.723 | 0.5 |  |
| Electron dose (e<sup>-</sup>/Å<sup>2</sup>) | 40 | 40 |  |
| Exposure rate (e<sup>-</sup>/Å<sup>2</sup>/sec) | 10 | 24 |  |
| Format of movies | EER | EER |  |
| Number of raw frames | 964 | 399 |  |
| Number of movies | 2,907 | 15,768 |  |
| Data processing |  |  |  |
| Number of fractions | 40 | 40 |  |
| Number of extracted particles | 391,755 | 1,369,874 |  |
| Number of particles for final map | 236,687 | 272,266 |  |
| Symmetry imposed | C1 | C1 | C1 |
| Resolution (Å) | 2.79 | 2.45 | 2.66 |
| FSC threshold | 0.143 | 0.143 | 0.143 |
| Refinement |  |  |  |
| Initial model used | 7MJG | 7MJM,7MJN | 7MJN |
| Map sharpening B-factor (Å<sup>2</sup>) | 72.2 | 39.2 | 82.6 |
| Composition (#) |  |  |  |
| Atoms | 21,960 | 33,398 | 6,567 |
| Residues | 2,703 | 4,087 | 796 |
| Ligands | NAG:58 | NAG:70 | NAG:7 |
| B-factor (Å<sup>2</sup>) |  |  |  |
| Protein (min/max/mean) | 45.69/247.52/111.11 | 25.51/351.21/165.88 | 52.33/188.61/88.85 |
| Ligand (min/max/mean) | 72.19/241.41/126.73 | 54.52/304.70/152.98 | 100.53/120.00/108.29 |
| Bonds (RMSD) |  |  |  |
| Length (Å) (# > 4σ) | 0.005 (0) | 0.004 (0) | 0.006 (0) |
| Angles (°) (# > 4σ) | 0.824 (8) | 0.842 (12) | 0.943 (4) |
| CC mask | 0.82 | 0.79 | 0.84 |

| **Validation** |  |  |  |
| --- | --- | --- | --- |
| Ramachandran plot |  |  |  |
| Residues favored (%) | 98.22 | 97.84 | 97.22 |
| Residues disallowed (%) | 0.04 | 0.02 | 0.00 |
| Rotamer outliers (%) | 0.17 | 0.03 | 0.14 |
| Clash score | 4.49 | 3.83 | 3.89 |
| MolProbity score | 1.23 | 1.21 | 1.32 |

**Table S2:**
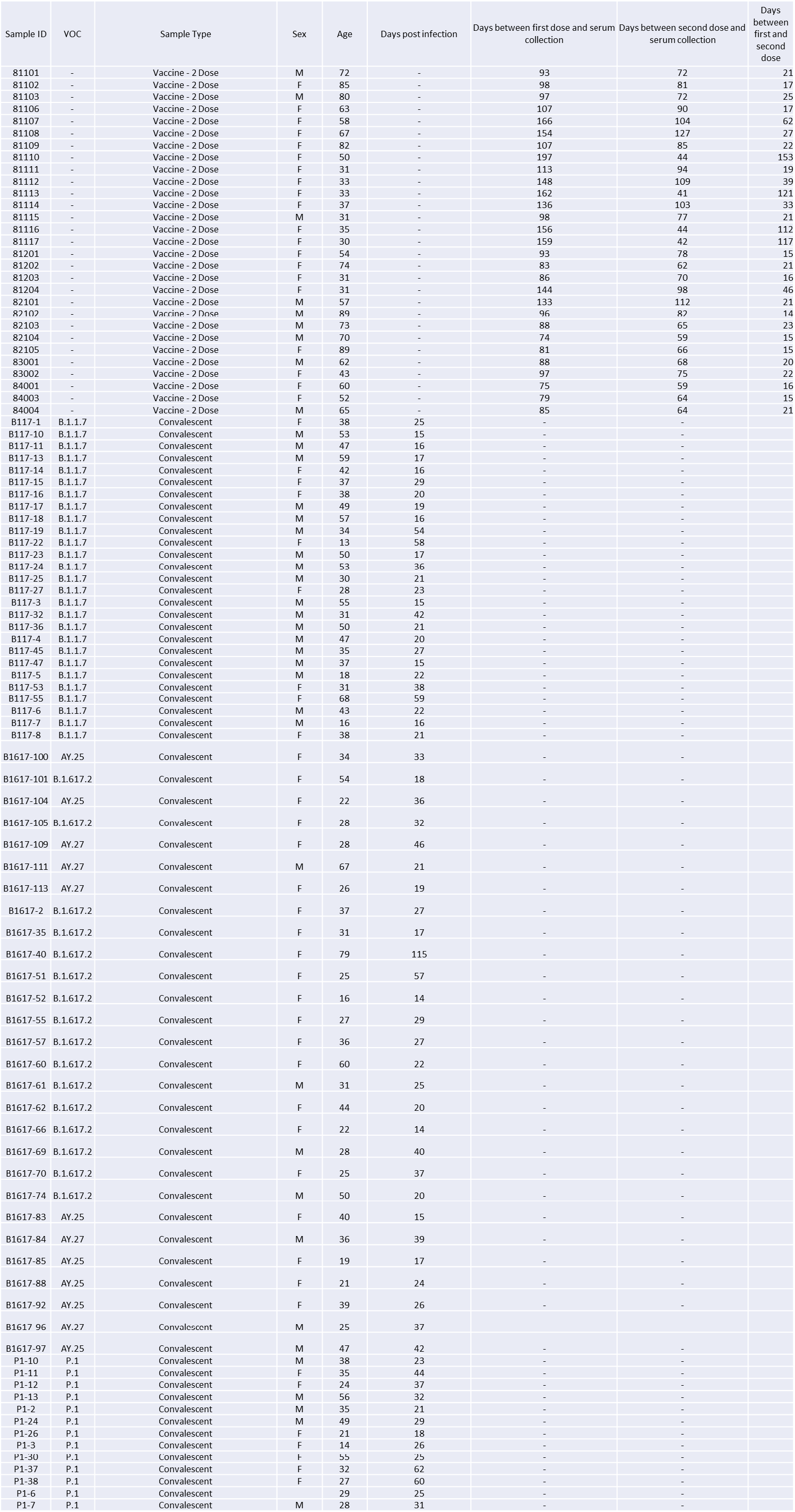
Summary of Patient Demographics

